# Hydraulic fracturing-induced delamination and extravasation extends medial damage beyond the false lumen in aortic dissection

**DOI:** 10.64898/2026.05.12.724712

**Authors:** Alan Chou, Abdulrahman H.M. Hassab, Jay D. Humphrey, George Tellides, Roland Assi

## Abstract

Aortic dissection is life-threatening due to continued loss of medial integrity that may culminate in secondary rupture within hours to days. While pre-existing defects or hemodynamic loads compound structural deterioration of the aorta, pathological progression from symptomatic dissection channel to lethal transmural tear is poorly understood. We examined the structure of referent and acutely dissected ascending aortas by microscopy. Elastic, collagen, and cellular components of non-dissected media were intricately interconnected. Medial damage in dissection lesions was traced from ingress to central to peripheral areas. Entry tears broke cleanly through successive laminae leading to cavernous false lumens in which medial structure was destroyed. Nearby laminae with widening between flanking elastic lamellae (termed minor delaminations) were filled with blood and showed severe medial damage. Farther laminae without delamination but containing red blood cells (termed blood extravasation) displayed moderate medial damage. More distant, non-delaminated laminae with accumulation of albumin but not red blood cells (termed plasma extravasation) exhibited mild medial damage. Varying medial hemorrhage with scattered sloughing of laminae was observed along the entire false lumen. We conclude that hydraulic fracturing of residual dissected media by pressurized blood via communications from the false lumen contributes to further structural weakening of the aortic wall.

## Introduction

The aorta is subject to substantial hemodynamic stress as it conveys blood from the heart to the body (1). Although the healthy aortic wall can withstand increased mechanical loads from strenuous activities, various conditions compromise structural strength of the dominant medial layer and render it vulnerable to failure. Loss of mural integrity includes tears through the intima and subjacent media that may: (i) heal partially or fully by fibrosis, forming a scar, (ii) split medial laminae in axial and circumferential planes, termed dissection, or (iii) extend radially through the entire thickness of the media, leading to either contained or free rupture (2). Healed intimomedial tears are benign, whereas ruptures are rapidly lethal, unless bleeding is contained by adventitia or perivascular tissue. Closer to the fatal end of the outcome spectrum, dissection of the proximal aorta has high mortality if untreated (∼3% immediately, ∼60% at 1 week, ∼75% at 2 weeks, ∼80% at 4 weeks, and ∼90% at 12 weeks), as documented in eras preceding cardiac surgical repair (3). Mortality is considerably higher for dissection of the ascending than descending thoracic aorta, even in the modern era (60% vs. 9% in-hospital mortality, respectively, in selected patients treated medically) (4). The vast majority of deaths (∼90%) in aortic dissection within 2 weeks of symptom onset are due to rupture with free bleeding into body cavities, mostly into the pericardial space (∼70%) because the ascending segment is most frequently involved (3). These considerations underscore an important pathological question of how dissection evolves to secondary rupture of the aortic wall.

Numerous post-mortem series during the past century and many contemporary histological studies of surgical specimens have focused on identifying structural defects predisposing to initial dissection. Almost all described an association with medial degeneration, namely elastin fragmentation, loss of smooth muscle cells (SMCs), accumulation of mucoid extracellular matrix (ECM), and fibrosis (3, 5–9). Yet medial degeneration is not found in all dissected aortas and is commonly associated with ageing in the absence of overt aortic disease (10, 11). Thus, medial degeneration is neither sufficient nor necessary for aortic dissection, although it likely plays a contributory role. In contrast to pre-existent disease, injury of the aortic media consequent to dissection has been infrequently studied. Since dissection channels typically track through the outer part of the media, the thinner layer of residual media external to the dissection channel is thought to underlie more frequent rupture of the ascending aorta outwards through the adventitia than re-entry tears inwards into the aortic lumen through the thicker inner layer of residual media and intima (3). Linear zones of SMC necrosis in the media internal to the dissection channel have been attributed to infarction from disruption of vasa vasorum that runs from adventitia to outer media of the thoracic aorta (12). This abnormality is observed in specimens from 48 hours to years after dissection occurrence, may ultimately cause collapse of acellular laminae though elastic lamellae remain intact, but is not documented as a nidus for further aortic tears. A few studies have characterized the evolution of inflammatory responses within the aortic wall from hours to months following dissection (13–15). The findings were related to reparative responses without assessment of medial damage; collections of blood were noted within the media adjacent to dissection channels, though of uncertain significance.

The present study was motivated by a need for improved understanding of aortic dissection complications. Mechanisms by which aortic dissection leads to secondary rupture are poorly understood, although the typical temporal delay from onset of symptoms to lethal outcome suggests that continued damage to the ascending aorta further weakens residual media and adventitia eventually leading to catastrophic structural failure. We find microscopy evidence of blood forced into medial laminae bordering the false lumen, separating adjacent elastic lamellae by breaking interconnections among elastic, collagen, and cellular components, thus resulting in varying and progressive medial damage. The pathological process is consistent with the theory of poroelasticity and engineering applications of hydraulic fracturing (16), and hydraulic fracturing of the aortic media represents a novel mode of disease pathogenesis.

## Results

### Interconnected ECM and cells of normal aortic media

To define normal medial structure, we examined ascending aortas from organ donors without aortic disease (*n* = 6, Supplemental Table 1 and Supplemental Figure 1). Consistent with prior observations (17), histology showed repeating units, termed laminae, of the dominant medial layer consisting of elastic lamellae bounding collagen fibers, glycoproteins, and SMCs (Figure 1A, B). Occasional small blood vessels, termed vasa vasorum, penetrated the outer media and interrupted the orderly layered organization. High-resolution confocal fluorescence microscopy confirmed that thin elastin protrusions, termed intralaminar elastic fibers, extended from thick elastic lamellae to SMCs with oblique alignment similar to cytoplasmic smooth muscle α-actin filaments (Figure 1C), known as the contractile-elastic unit (18). The fibrillar collagen matrix comprised predominantly type I and III collagen fibers arranged parallel to elastic lamellae and extending between SMCs (Figure 1D, E). Type IV collagen, a basement membrane component, surrounded SMCs contributing to a narrow, uneven pericellular matrix distinct from the fibrillar collagen matrix and extended along neighboring cells (Figure 1E), known as the pericellular binding matrix (19). An irregular cell membrane delineated SMC projections that attached to intralaminar elastic fibers with greater density of integrin receptors at points of contact (Figure 1F), likely reflecting focal adhesions (20). Hence, medial laminae have extensive connections among the elastin framework, collagen fibers, basement membrane, and cells.

**Figure 1:**
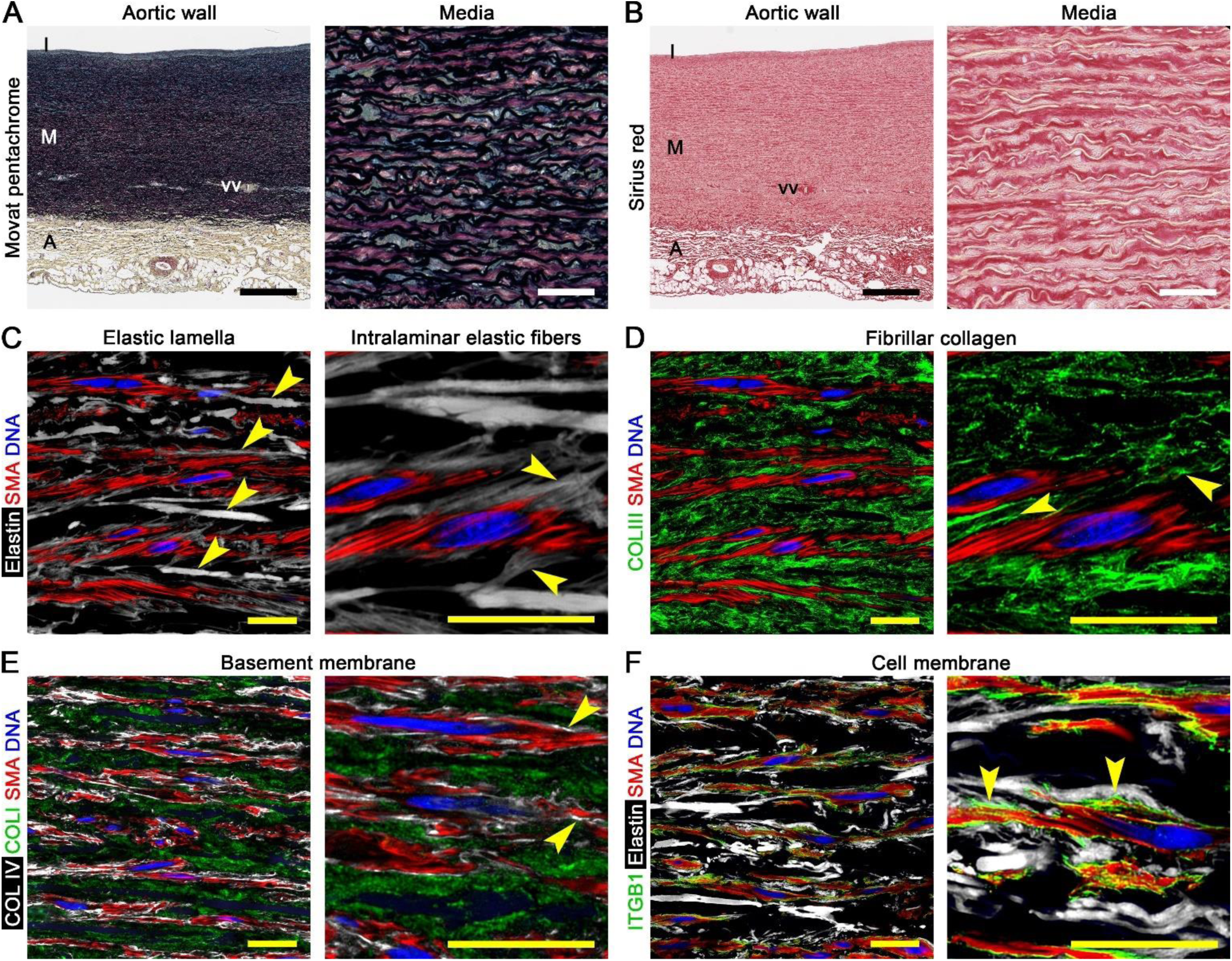
Interconnected ECM and SMCs of normal aortic media. Histology and confocal fluorescence microscopy images of non-diseased aortas from organ donors. (**A**) Movat pentachrome stain labeling elastin (black), cells (red), collagen (yellow), and glycosaminoglycans (blue) delineating the intima (I), media (M), and adventitia (A) with penetrating vasa vasorum (vv). (**B**) Sirius red stain labeling collagen (red). (**C**) Hydrazide-labeled elastin (white) revealing parallel, thick elastic lamellae (arrows, left panel) with extensions of oblique, thin intralaminar elastic fibers (arrows, right panel) attached to SMCs that contain similarly oriented filaments of smooth muscle α-actin (SMA, red) and DAPI-labeled nucleic DNA (blue). (**D**) Type III collagen (COLIII, green) delineating collagen fibers (arrows, right panel) adjacent to elastic lamellae and between SMCs. (**E**) Type IV collagen (COLIV, white) delineating basement membrane surrounding SMCs as thin pericellular matrix (arrows, right panel) separate from type I collagen (COLI, green) of fibrillar collagen matrix. (**F**) Integrin-β1 (ITGB1, green) outlining irregular cell membrane of SMCs with greater density (arrows, right panel) at attachments to intralaminar elastic fibers. Microscopic fields: normal media at varying magnification to highlight various structures (panels A–F). Orientation: luminal side top, adventitial side bottom. Scale bars: black = 500 μm, white = 50 μm, yellow = 20 μm.

### Entry tears break successive laminae with minimal damage to adjacent inner media

To characterize medial damage from ascending aortic dissection, we examined excised specimens from patients undergoing emergent surgical treatment within 24 hours of symptom onset (*n* = 11, Supplemental Table 2 and Supplemental Figure 1). Medial structure was examined systematically by tracing tissue injury along a sequence of pathological events that enable blood to enter the vessel wall from the lumen. Initial tears through the intima and subjacent media, known as entry tears, were readily apparent on gross inspection with varying appearances (Supplemental Figure 2A–E). By histology, entry tears had a corrugated appearance and broke multiple elastic lamellae in a radial direction terminating in a cavernous aberrant channel, known as the false lumen, deep within the media (Figure 2A). The inner media adjacent to entry tears did not show obvious damage, having an orderly parallel arrangement of elastin and collagen similar to intact media not affected by dissection in the same specimens. Confocal fluorescence microscopy of the inner media bordering entry tears confirmed absent blood extravasation and relatively undamaged elastic lamellae, intralaminar elastic fibers, collagen fibers, basement membrane, and SMCs (Figure 2B, C). In some dissected aortas, sparing of the inner media adjacent to entry tears was confounded by antecedent medial degeneration (Supplemental Figure 3A–C). Thus, entry tears represent focal intimomedial disruptions in compromised aortas but without apparent secondary injury of the adjacent inner media.

**Figure 2:**
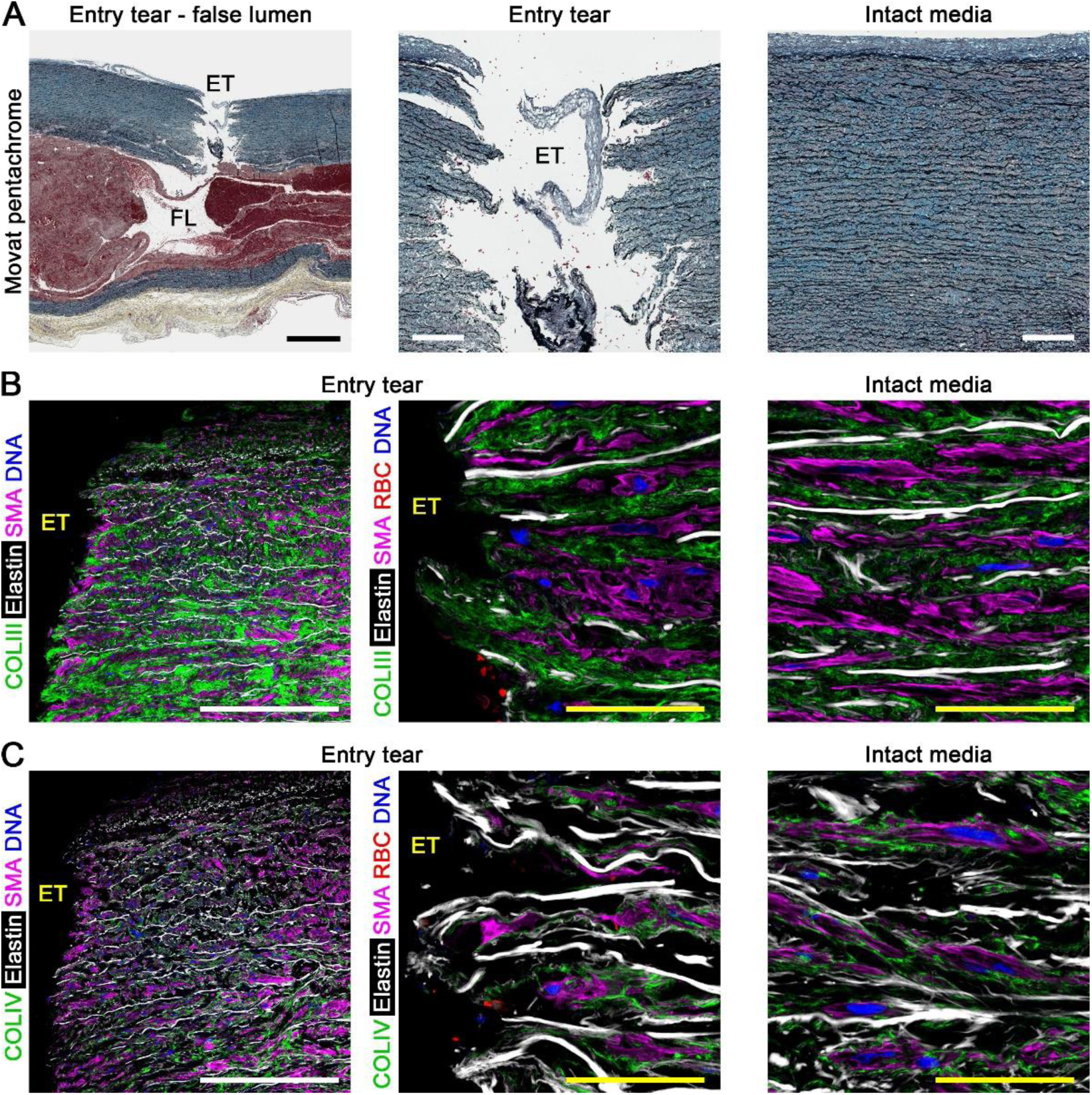
Entry tears break successive laminae with minimal damage to adjacent inner media. Histology and confocal fluorescence microscopy images of dissected aortas focused on entry tears compared to intact (non-dissected) media in the same subject. (**A**) Movat pentachrome stain labeling elastin (black), cells and fibrin (red), collagen (yellow), and glycosaminoglycans (blue) showing an entry tear (ET) opening into a false lumen (FL). (**B**) Collagen III (COLIII, green) delineating collagen fibers, hydrazide labeling elastin (white), smooth muscle α-actin (SMA, magenta) identifying SMCs, CD235a (red) identifying RBCs, and DAPI labeling DNA of nuclei (blue). (**C**) Collagen IV (COLIV, green) delineating basement membrane that surrounds SMCs. There is no blood extravasation and little damage of the elastin framework or fibrillar collagen and basement membrane matrix in media adjacent to entry tears versus intact media. Microscopic fields: entry tear and false lumen (panel A left), entry tear (panels A center, B left and center, C left and center), intact media (panels A right, B right, C right). Orientation: luminal side top, adventitial side bottom. Scale bars: black = 1,000 μm, white = 250 μm, yellow = 50 μm.

### Destruction of media within markedly separated laminae of false lumen

We examined the false lumen that extends axially and circumferentially from the entry tears. False lumens were filled with blood and thrombus, ranging from several hundred μm to several mm or cm in radial dimension, and were delimited by residual media internally (termed inner media or dissection flap) and externally (termed outer media), compressing the contiguous true lumen (Supplemental Figure 2F). Some false lumens were complex with compartments communicating out of the plane of section. The interior of the false lumen was devoid of ECM and SMCs, except debris (Figure 3A). Occasional sheets of fractured media lining the false lumen sloughed into the dissection cavity (Figure 3B). The cavity edge irregularly traversed laminae across breaks in contiguous elastic lamellae (Figure 3C). Most exposed elastic lamellae were stripped clean of intralaminar elastic fibers, collagen fibers, basement membrane, and cells (Figure 3D, E). Loss of intralaminar elastic fibers was also evident in laminae bordering the false lumen whether flanking elastic lamellae were separated or not, whereas fibrillar and pericellular collagen were relatively intact in non-delaminated laminae. The magnitude of tissue injury indicates complete destruction of medial structure within the widely separated laminae contributing to the false lumen due either to intramural stress initially splitting the media and/or continued inflow of blood.

**Figure 3:**
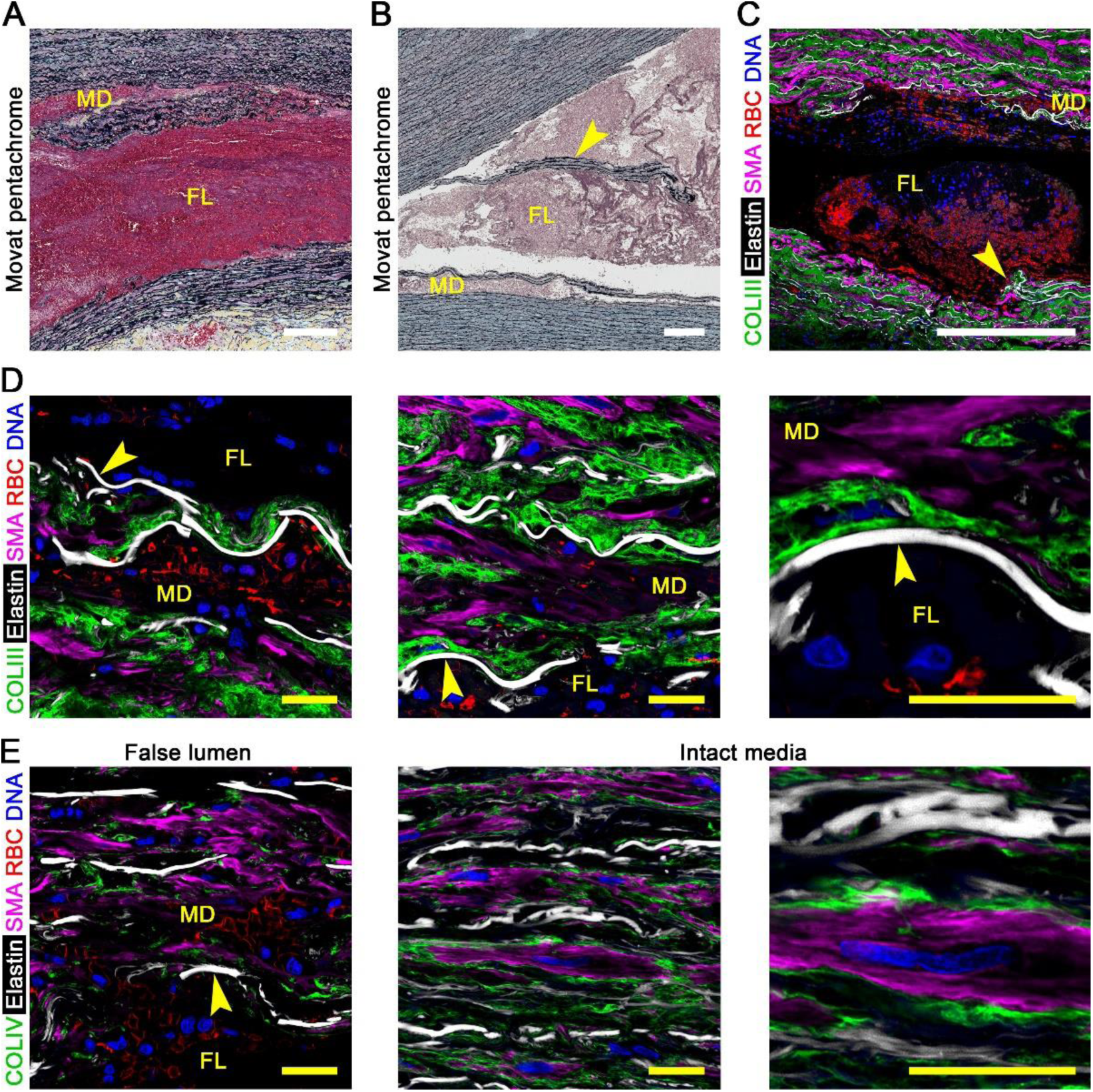
Medial destruction in widely separated laminae forming false lumen. Histology and confocal fluorescence microscopy images of dissected aortas focused on false lumen compared to intact media in the same subject. (**A**) Movat pentachrome stain showing false lumen (FL) and nearby minor delamination (MD) containing RBCs and fibrin (red). (**B**) In some sections, sheets of medial laminae (arrow) had dislodged into the false lumen. (**C**) Collagen III (COLIII, green) delineating collagen fibers, hydrazide (white) labeling elastin, smooth muscle α-actin (SMA, magenta) identifying SMCs, CD235a (red) identifying RBCs, and DAPI (blue) labeling nucleic DNA with contiguous breaks in elastic lamellae (arrow) permitting the cavity edge to cross laminae. (**D**) Higher magnification views of exposed elastic lamellae bordering the false lumen stripped clean of fibrillar collagen and cells (arrows). (**E**) Collagen IV (COLIV, green) delineating basement membrane is also stripped clean from elastic lamellae exposed in the false lumen (arrow) but surround SMCs in intact media. Microscopic fields: false lumen (panels A–D, E left), intact media (panels E center and right). Orientation: luminal side top, adventitial side bottom. Scale bars: white = 250 μm, yellow = 20 μm.

### Severe medial damage in laminae with minor delamination

We examined widened laminae that contained blood, near to but distinct from the false lumen, referred to as minor delaminations (the false lumen is considered the major delamination). In the absence of dissection, the thickness of individual medial laminae in non-pressurized formalin-fixed sections, not including flanking elastic lamellae, is 15.6 ± 3.3 μm in non-dilated aortas and 16.5 ± 5.7 μm in dilated aortas (21). Minor delaminations ranged from 25–100 μm in width, sometimes single, sometimes several in parallel, and were filled with red blood cells (RBCs) visualized via different histological stains (Figure 4A–C). Most minor delaminations were within 5–10 laminae from the border of the false lumen, though clear communications with the false lumen were seldom. Minor delamination was also observed just distal to the end of the false lumen, termed the dissection front. The interior of minor delaminations was devoid of ECM and SMCs, except for debris. At the edges, elastic lamellae were often stripped clean with occasional fragments of collagen and SMCs attached (Figure 4D, E). Few platelets were detected in minor delaminations, although abundant within the false lumen (Supplemental Figure 4A, B). Thus, separation of elastic lamellae with consequent traction on attached medial components and/or intralaminar blood flow results in extensive connective tissue and cellular damage within minor delaminations as in the false lumen.

**Figure 4:**
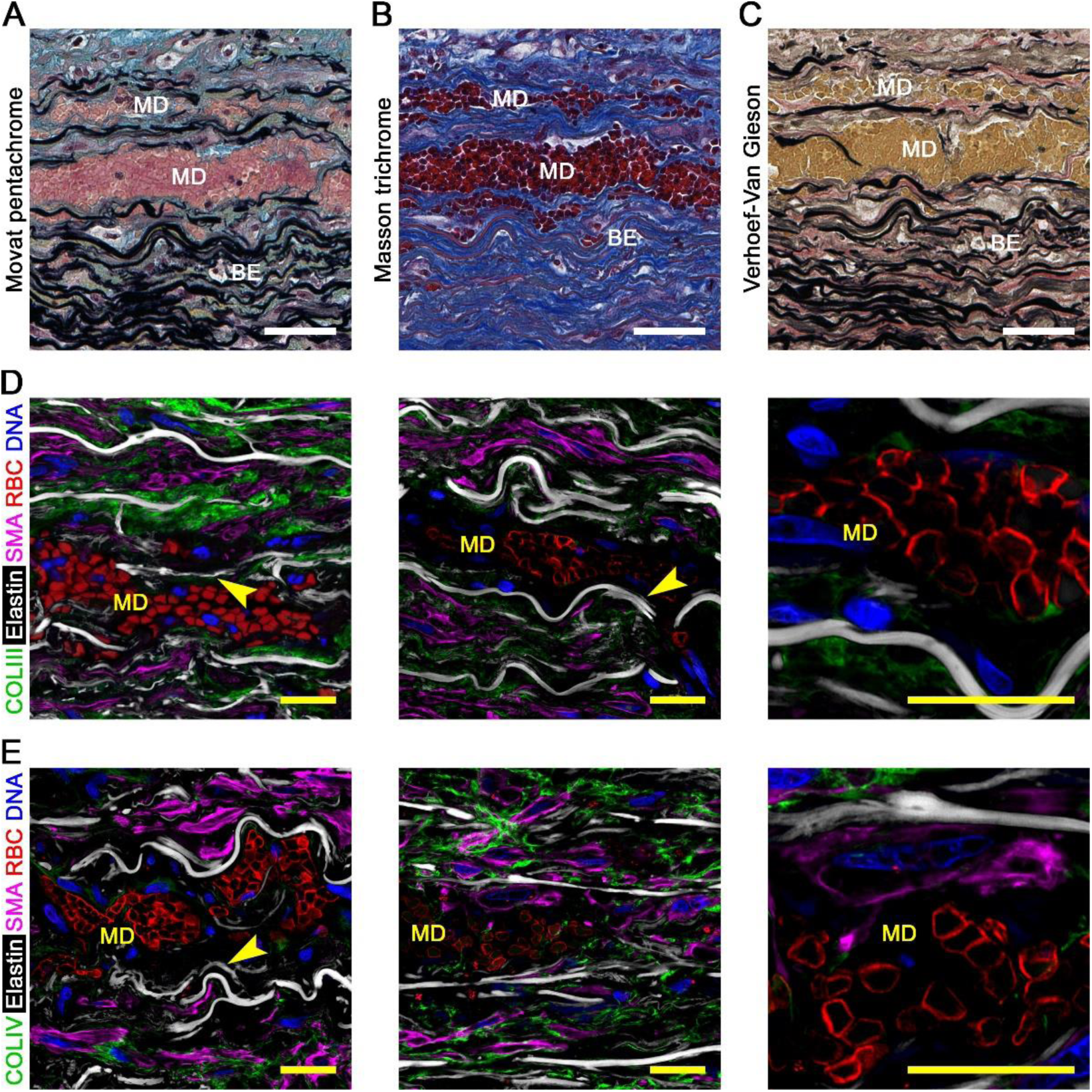
Severe medial damage in laminae with minor delamination and blood extravasation. Histology and confocal fluorescence microscopy images of dissected aortas focused on laminae with minor delaminations distinct from false channel. (**A**) Movat pentachrome stain showing laminae with minor delamination (MD) filled with blood and thrombus (pink-red) in addition to adjacent, non-widened laminae containing RBCs, termed blood extravasation (BE). (**B**) Masson trichome stain identifying RBCs (dark red) among layers of collagen fibers (blue). (**C**) Verhoeff-Van Gieson stain identifying RBCs (yellow) among layers of elastic fibers (black). (**D**) Collagen III (COLIII, green) delineating collagen fibers, hydrazide (white) labeling elastin, smooth muscle α-actin (SMA, magenta) identifying SMCs, CD235a (red) identifying RBCs, and DAPI (blue) labeling nuclei, with (**E**) collagen IV (COLIV, green) instead of collagen III showing severe damage to medial components in areas of minor delamination with parts of exposed elastic lamellae stripped clean of fibrillar collagen and cells (arrows). Microscopic fields: minor delaminations (panels A–E). Orientation: luminal side top, adventitial side bottom. Scale bars: white = 50 μm, yellow = 20 μm.

### Moderate medial damage in non-delaminated laminae with blood extravasation

We examined laminae containing blood but without appreciable separation of flanking elastic lamellae, referred to as blood extravasation, mostly situated farther from the false lumen or more distal to the dissection front than laminae with minor delamination. The width of non-delaminated laminae was < 25 μm with fewer RBCs than in minor delaminations. The outlines of RBCs interfaced with distorted and ruptured SMCs (Figure 5A, B). Furthermore, RBCs were detected between elastic lamellae and collagen fibers, between collagen fibers and basement membrane, between basement membrane and SMCs, and within ruptured SMCs. High magnification confocal images demonstrated RBCs protruding into disruptions in the basement membrane or cell membrane of SMCs (Figure 5C, D). Minimal fibrin was detected in non-widened laminae with blood extravasation, though conspicuous in minor delaminations and the false lumen (Supplemental Figure 5A, B). These findings suggest that extravasation of blood through myriad small communications from the false lumen disrupts interconnections among ECM and cells, fragments collagen, and ruptures SMCs, even in the absence of tractions from elastic laminae pulling apart suggesting direct injury from pressure and/or flow.

**Figure 5:**
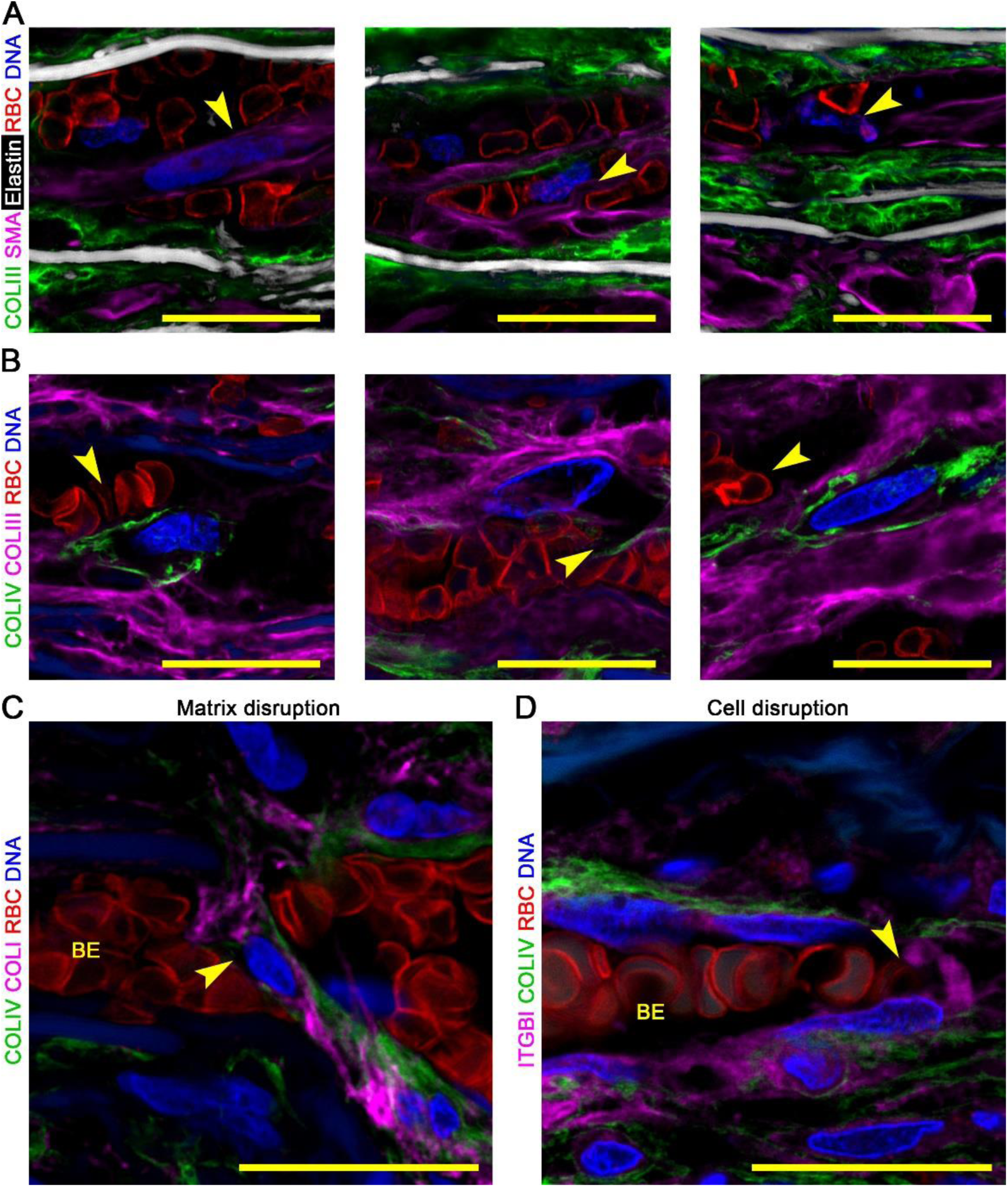
Moderate medial damage in non-delaminated laminae with blood extravasation. Confocal fluorescence microscopy images of dissected aortas focused on non-widened laminae with blood extravasation (BE). (**A**) Collagen III (COLIII, green) delineating collagen fibers, hydrazide (white) labeling elastin, smooth muscle α-actin (SMA, magenta) identifying SMCs, CD235a (red) identifying RBCs, and DAPI (blue) labeling nuclei showing disruption of perinuclear cytoplasm (arrows). (**B**) Collagen IV (COLIV, green) delineating basement membrane, collagen III (magenta), CD235a (red), and DAPI (blue) showing blood extravasation between basement membrane and collagen fibers (arrow, left panel), between basement membrane and cell (arrow, middle panel), and between collagen fibers (arrow, right panel). (**C**) Collagen IV (green), collagen I (COLI, magenta) delineating collagen fibers, CD235a (red), and DAPI (blue) showing discontinuity in basement membrane (arrow). (**D**) Collagen IV (green), β1 integrin (ITGB1, magenta) delineating cell membrane, CD235a (red), and DAPI (blue) showing discontinuity and peeling of cell membrane (arrow). Elastin is barely visible by autofluorescence (dull blue) in the absence of hydrazide (panels B–D). Microscopic fields: blood extravasation (panels A–D) with high magnification views of ECM disruption (panel C) and cell disruption (panel D). Orientation: luminal side top, adventitial side bottom. Scale bars: yellow = 20 μm.

### Mild medial damage in non-delaminated laminae with plasma extravasation

We examined laminae with neither separation of elastic lamellae nor infiltration with RBCs in lesion areas distant from the false lumen. Plasma infiltration was inferred by dense accumulation of albumin, although this parameter could not be reliably applied to all specimens as the typical patchy distribution of albumin was more intense within the intact media of some dissected aortas and even in parts of the media of non-dissected aortas suggesting breakdown of usual endothelial or elastic laminae permeability barriers (Supplemental Figure 6A–D). In aortic dissection specimens without increased albumin in the intact media, there was a central zone of predominantly RBCs with less albumin and no SMCs corresponding to the false lumen and minor delaminations, a transitional zone of comparable RBCs and albumin corresponding to blood extravasation with few residual SMCs, and a peripheral zone of albumin with many SMCs but no RBCs, termed plasma extravasation (Figure 6A, B). In non-delaminated laminae with plasma extravasation, albumin accumulation was largely pericellular and between elastic lamellae and collagen fibers (Figure 6C). In some lesion areas, albumin accumulated within SMCs but did not localize to sparse lysosomes (Figure 6D, E); detection of lysosomal membrane proteins was confirmed in phagocytic leukocytes frequent in false lumens and minor delaminations but absent in plasma extravasation (Supplemental Figure 7A–E). Distortion of SMCs in lesion areas with dense albumin accumulation was indicated by altered orientation of nuclei (Figure 6F). In rare areas of plasma extravasation, SMCs were disrupted with cytoplasm separated from isolated nuclei that were surrounded by albumin (Figure 6G). Accordingly, intramural fluid flow, through sufficiently small channels that restrict blood cells but not plasma proteins, is associated with sporadic medial damage.

**Figure 6:**
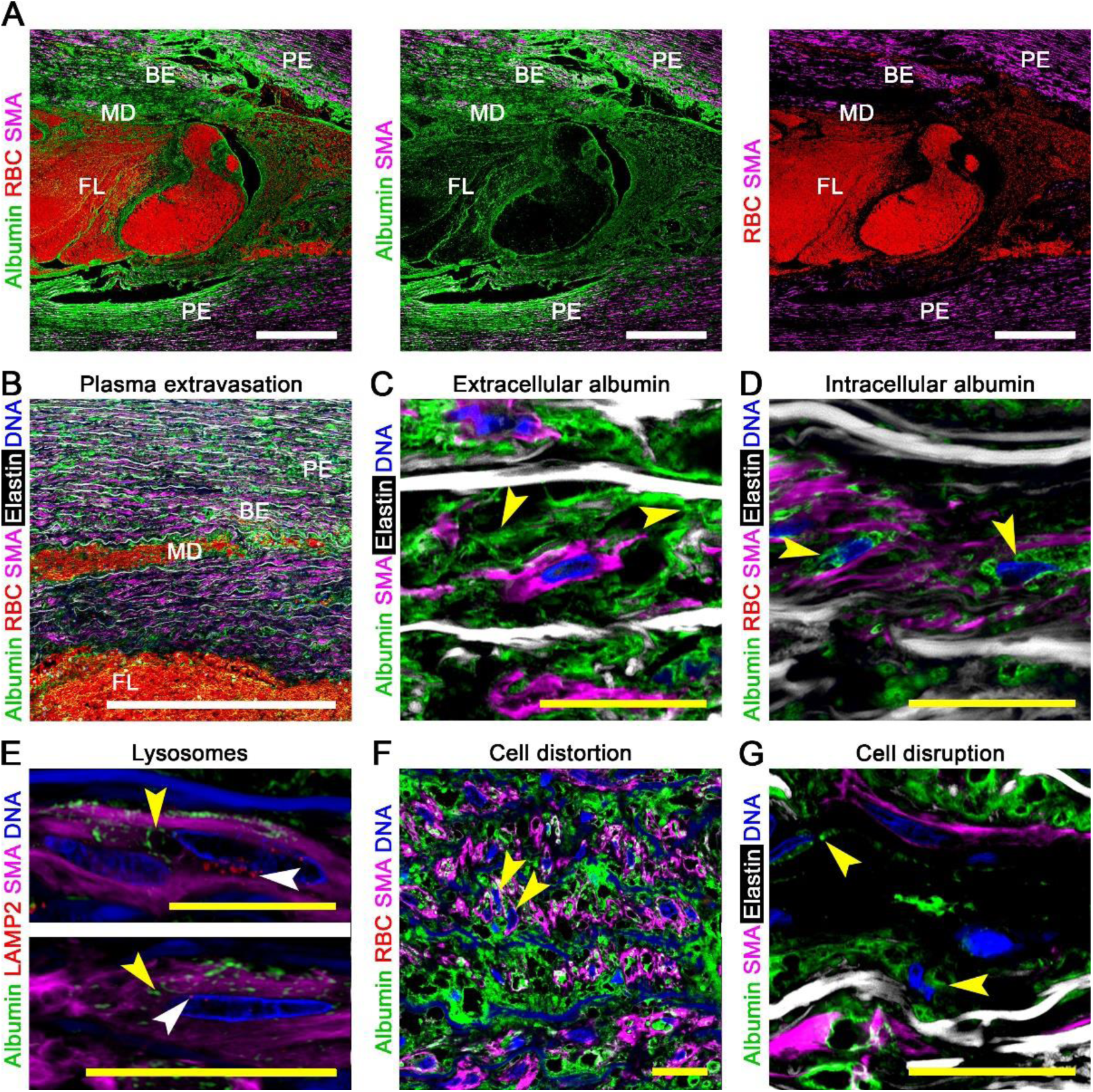
Mild medial damage in non-delaminated laminae with plasma extravasation. Confocal fluorescence microscopy images of dissected aortas focusing on laminae without separation of elastic lamellae or RBCs but with plasma protein infiltration, termed plasma extravasation. (**A**) Varying infiltration of albumin (green) and RBCs identified by CD235a (red) among smooth muscle α-actin (SMA, magenta) containing SMCs showing a central false lumen (FL), adjoining zones of medial delamination (MD) and blood extravasation (BE), and peripheral zones of plasma extravasation (PE). (**B**) Low magnification view of albumin (green), RBCs (red), SMCs (magenta), elastin (white), and DAPI-labelled nuclei (blue) identifying a hierarchy of medial injury. (**C**) Medial albumin accumulation after dissection is often extracellular, though (**D**) intracellular in some areas. (**E**) Albumin (green, yellow arrows) that accumulated internal to smooth muscle α-actin filaments (magenta) did not localize to perinuclear lysosomes delineated by lysosomal-associated membrane protein 2 (LAMP2, red, white arrows). (**F**) Altered orientation of nuclei (arrows) within areas of dense albumin accumulation indicating distortion of SMCs. (**G**) Uncommonly, albumin (green, yellow arrows) encircled isolated nuclei (blue) stripped of cytoplasm (magenta). Microscopic fields: aortic dissection lesion (panels A), plasma extravasation (panel B), extracellular albumin accumulation (panels C), intracellular albumin accumulation (panel D), albumin and lysosome distribution (panel E), distorted SMCs (panel F), and disrupted SMCs (panel G). Orientation: luminal side top, adventitial side bottom. Scale bars: white = 500 μm, yellow = 20 μm.

### Widespread medial hemorrhage along the false lumen

We characterized the extent of medial injury through surveys of low magnification histophotomicrographs of dissected aortas. Detection of RBCs within the media adjacent to the false lumen is consistent with either minor delaminations or blood extravasation (can be difficult to distinguish on histological stains that do not discriminate elastic lamellae) and is collectively referred to as medial hemorrhage. A similar extent of hemorrhage was noted within 5–10 laminae adjacent to the false lumen in both inner and outer medial layers (Figure 7A, B). Of 11 subjects examined, 6 had specimens of both proximal and distal ascending aorta representing areas of varying distance from the entry tear; there was little difference in the extent of medial hemorrhage with respect to distance from the entry tear (Figure 7C, D). In specimens with severe medial degeneration, substantial hemorrhage occasionally extended into areas of proteoglycan-rich, cell-and elastin-deficient media appearing as secondary blood channels distinct from the primary false lumen (Figure 7E). In comparison, bleeding into media with mild degeneration was more limited (Figure 7F). Typically, more extensive lesions were seen near the dissection front where a hemorrhagic network spread up to 1 mm (equivalent to ∼50 laminae) from the border of the false lumen (Figure 7G). Rarely, radial tears extended across the outer media distant from the entry tear allowing blood to reach the subjacent adventitia illustrating how aortic rupture can subsequently occur (Figure 7H). These findings reveal bleeding into the media along the length of the false channel thus enabling further medial damage and possible secondary rupture at any point within the dissected aorta.

**Figure 7:**
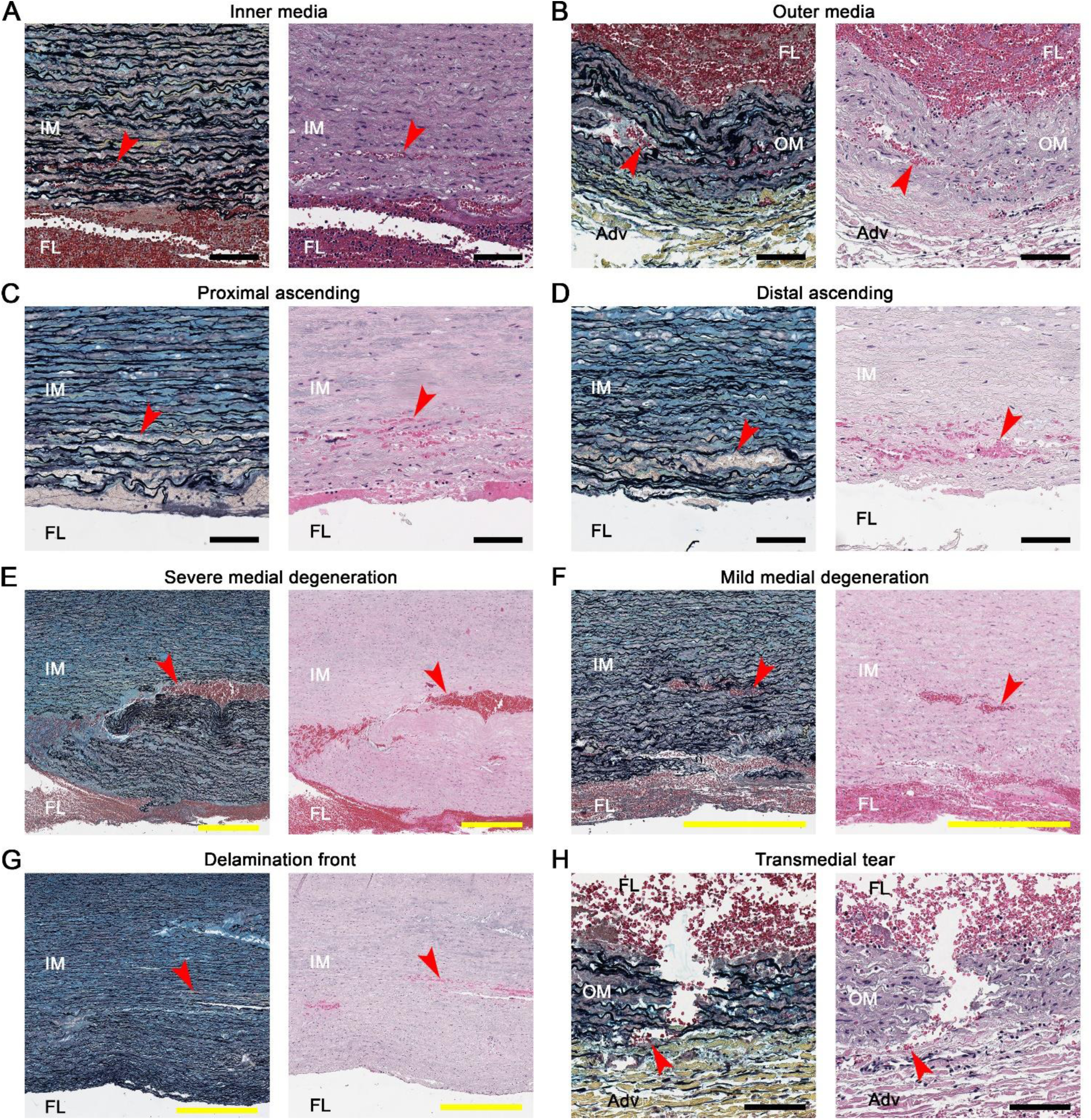
Widespread medial hemorrhage along the false lumen. Histophotomicrographs of aortic dissections demonstrating the extent of medial hemorrhage (comprising minor delamination and blood extravasation lesions). Movat pentachrome stains (left panels) and hematoxylin and eosin stains (right panels) label RBCs a red or pink (arrows). Similar extent of medial hemorrhage in (**A**) inner media (IM) and (**B**) outer media (OM) to the false lumen (FL), and in (**C**) proximal and (**D**) distal ascending aorta, regardless of distance from the entry tear. (**E**) Substantial hemorrhagic channels (100–200 µm width) form in areas with severe medial degeneration, contrasted with (**F**) more limited hemorrhage in media with mild degeneration. (**G**) Greater extent of medial hemorrhage (up to 1 mm) near the delamination front of the false lumen. (**H**) Radial tear extending through the outer media with blood reaching the adventitia (Adv). Scale bars: yellow = 500 μm, black = 100 μm.

## Discussion

We document a decreasing outward gradient of damage to the media surrounding the false lumen in aortic dissection comprising: (i) minor delaminations of widened laminae filled with blood in the absence of ECM and SMCs, except for occasional fragments adherent to lining elastic lamellae, (ii) non-delaminated laminae with extravasation of RBCs that show destruction or displacement of intralaminar elastic fibers, collagen fibers, basement membrane, and SMCs, and (iii) non-delaminated laminae with extravasation of albumin but not RBCs that exhibit infrequent ECM and SMC damage. These observations implicate successive modes of medial injury from fluid flow under pressure (extravasation) and traction on ECM and SMCs firmly attached to flanking elastic lamellae that are forced apart (delamination). Persistent attachment of SMC fragments to elastic laminae indicates that these cell membrane connections are not dependent on cell viability. Separation of SMCs from adjacent collagen fibers and basement membrane by extravasated blood demonstrates loss of structural connections among elements of the pericellular binding matrix; SMC damage was not evident in the presence of an intact basement membrane. Loss of structural integrity of the media external or internal to the false lumen within dissected aortas may result in transmedial tears (that may be contained by adventitia and perivascular tissue), transmural tears (with lethal consequences if there is free bleeding into a body cavity), or re-entry tears (that reestablishes blood flow into the true lumen).

Medial constituents display varying susceptibility to injury along lesions. Elastic lamellae sustain multiple successive breaks in entry tears, show irregular contiguous breaks within the false lumen that allow cavity edges to cross laminae, but are relatively intact in peripheral lesion areas with extravasation. In contrast, intralaminar elastic fibers are more ephemeral and often not detected in laminae (both with and without delamination) close to false lumens, perhaps breaking from marked distortion of the media associated with rapid enlargement of dissection channels. Type I and III collagen fibers, largely parallel to elastic lamellae, and type IV collagen, around SMCs, are absent in minor delaminations, fragmented, peeled, or distorted in lesion areas with blood extravasation, and occasionally damaged in lesion areas with plasma extravasation. Strips of basement membrane separated from the surface of SMCs were previously reported in rabbit aortas subjected to hyperdistention from elevated pressures ex vivo (19). SMCs follow a similar pattern of injury to ECM, also manifesting intracellular albumin accumulation in some lesion areas with plasma extravasation. The mechanism for albumin entry into cells is unknown and may occur from receptor-mediated endocytosis (22) or non-specific, actin-dependent macropinocytosis (23, 24), although there is no evidence for eventual trafficking to the endosomal-lysosomal compartment. Alternatively, albumin entry may occur via plasma membrane wounds secondary to hydraulic forces that are rapidly repaired from a lysosomal pool of membrane (25), but lysosomal membrane protein is not detected at the cell surface. Increased fluid transport through the residual dissected aortic wall agrees with intraoperative observations that fluid can seep through the thin outer media and adventitia resulting in pericardial effusions (26) and electrocautery is less effective in separating dissected tissues, as if saturated with fluid (27).

Hydraulic fracturing is an engineering term that refers to the creation of fissures in porous materials of low permeability in which a pressurized liquid is forced through confined spaces. Also known as hydrofracturing and in the vernacular as fracking, the concept underlies the reservoir stimulation process to maximize extraction of underground resources by the oil and natural gas industry. The procedure involves drilling vertical wells with horizontal shafts and pumping in water containing additives at high pressure that fractures surrounding deep rock formations; once pumping ceases, a proportion of fracturing fluid bearing hydrocarbons flows back to the surface (28). As in geotechnical applications, hydraulic fracturing based on differences in fluid flow and pressure have also been described in physiological cell functions within the theoretical framework of poroelasticity (29–31). Examples include detachment of cell membrane from the underlying cytoskeleton during blebbing (32, 33), breaks in nuclear envelopes during cell migration through confining environments (34, 35), and disruption of adhesive contacts separating cells (36, 37). While hydraulic fracturing is not specifically invoked in pathogenesis, high-pressure injection injuries from industrial tools in the workplace can result in digit amputations (38, 39) and numerous case reports describe soft tissue injuries in children from high-pressure washer devices (40–42), even water/air infiltration from a standard pressure bath faucet in the context of penetrating trauma (43). Conversely, hydraulic fracturing has therapeutic applications for separating tissues with less trauma than cutting instruments by a technique of hydrodissection using directed jets of fluid (44).

Principles of hydraulic fracturing from rock mechanics may inform studies of aortic dissection. Entry tears and false lumens are analogous to vertical and horizontal wellbores, while cardiac contractility generates cyclical blood pressure allowing antegrade and some retrograde motion of fluid. Fissures in the media on either side of the false lumen maintain tortuous communications with the circulation. The orientation of most fracture channels is parallel to elastic lamellae, and their size generally decreases from central to peripheral lesion areas. In comparison to the normal vasculature, false lumens can exceed aortic true lumen diameter (several cm), minor delaminations are about the width of arterioles (25–100 μm), blood extravasation channels are around capillary size (5–25 μm), and plasma extravasation channels are likely smaller than capillaries since they restrict RBC traffic. While pressures were not determined, they predictably diminish along the direction of flow (as in the arterial tree from larger to smaller vessels), and thrombus within the false lumen or minor delaminations may further decrease luminal pressure. The contents of medial fractures are related to their size; RBCs are absent in plasma extravasation channels and leukocytes are infrequent in blood extravasation channels. Blood is unlikely to enter the residual media through fenestrations of elastic laminae lining the false lumen as these are typically 0.5–1.5 μm in size in the thoracic aorta (45), though may enlarge with aortic disease. Blood has greater viscosity than plasma (46) which influences fracturing abilities. Albeit not considered herein, the state of biaxial wall stress is also expected to influence the fracture mechanics (47). Biological processes further modulate channel contents. Coagulation reactions entrap platelets and/or fibrin within the false lumen and minor delaminations but are not detected within extravasation areas. Thrombus may also release and activate proteases contributing to mural degradation (48). Furthermore, passive and active material properties of the aortic wall, in particular weakening of medial ECM and loss of SMC contractility, impact vulnerability to structural failure (49). Of relevance, pressure (including cyclic variation), viscosity, permeability, porosity, and mechanical properties are determinants of hydraulic fracture channels in geotechnical applications (28, 50), supporting applicability of these variables to fracturing of the aortic media.

Our studies in human specimens were inspired by findings in mouse models of aortopathy. Unlike classic aortic dissection, medial hemorrhage in murine ascending aortas uncommonly results in false lumens—instead blood frequently infiltrates multiple laminae (51, 52). Analysis of acute dissection lesions within 30 minutes of increasing blood pressure in adult mice with disrupted TGF-β signaling in SMCs reveals tortuous communications from entry tears to fracture channels with blood extravasation, varying separation of adjacent elastic lamellae, and progression from circumferentially to radially oriented SMCs to cell body rupture with increasing delamination (53). Widened laminae were filled with RBCs and lining elastic lamellae contained fragments of SMCs and collagen similar to minor delamination and blood extravasation lesions described herein. In contrast, aortic tears in descending thoracic and suprarenal abdominal segments in mice often manifest as contained ruptures, though pharmacological inhibition of mTOR signaling converts some lesions to medial extravasation and delamination characteristic of dissection while markedly decreasing the overall incidence of transmedial tears (54). In human aortic dissection, tears extending through the outer media resulting in rupture are more common in the ascending segment, while tears extending through the inner media resulting in re-entry to the true lumen are more common in descending thoracic and abdominal segments (3, 55). Other than the aforementioned study in which rapamycin did not prevent production of matricellular proteins or modulation of SMC adhesion induced by high-dose angiotensin II infusion in mice (54), there is little insight into mechanisms that differentially regulate radial tears versus axial/circumferential delamination of the aortic wall.

Hydraulic fracturing of the media in aortic dissection adds further complexity to our recent description of a multiphasic model of aortic wall failure (2). This concept emphasizes the resilience of healthy aortas in which intimomedial tears remain limited and heal by SMC-mediated scar formation versus the vulnerability of diseased aortas in which intimomedial tears (entry tears in the context of aortic dissection) extend as dissection or further complicate as rupture. The current work highlights a network of fracture channels connected to the false lumen and illustrates a mechanism for progressive damage of residual media (e.g., cleavage of sheets of laminae shown in Figure 3B or radial tear extension shown in Figure 7H) that may contribute to rupture. While our observations are informative about a novel mode of pathology and the cellular and extracellular components affected, further mechanistic studies are required to define biological processes and molecular functions involved to identify potential therapeutic targets. Hydraulic fracturing may apply to other diseases driven by pressurized fluid, such as atherosclerotic plaque hemorrhage and rupture.

## Methods

### Study approval

Research protocols to obtain aortic tissue from surgical patients were approved by the Yale Institutional Review Board with a waiver for consent. Research protocols to obtain aortic tissue from deceased organ donors were approved by the Yale Institutional Review Board with waiver of consent and by the New England Organ Bank with written consent for research from the next-of-kin. All procedures were in accordance with federal and institutional guidelines.

### Subjects

Ascending aorta specimens were collected from October 2021 to December 2023 from 11 subjects who underwent emergent surgery for acute aortic dissection and from October 2017 to February 2022 from 6 organ donors whose hearts were not used for transplantation. The specimens were procured by the investigators in the operating room to ensure correct anatomical location and orientation. Duration of aortic dissection (9.7 ± 8.2 hr, range: 3–24 hr) was calculated from the onset of symptoms to the time of aortic cross-clamping shortly before aortic resection.

### Histology

Transverse sections of non-pressurized aortas were fixed in 4% paraformaldehyde overnight at 4 °C and embedded in paraffin. Tissue blocks were sectioned at 5 μm thickness and stained with Movat’s pentachrome, Masson’s trichrome, Verhoef-Van Gieson, and sirius red stains by Yale’s Research Histology Laboratory using standard techniques. Slides were digitized using an Aperio AT2 scanner (Leica).

### Confocal fluorescence microscopy

Similar formalin-fixed, paraffin-embedded aortic specimen slides were serially dewaxed in xylene and rehydrated in a series of graded alcohol then water. Heat-mediated antigen retrieval was performed (H-3300-250, Vector Laboratories). Sections were incubated overnight at 4 °C with antibodies to smooth muscle α-actin (clone 1A4, mouse IgG2a, Alexa Fluor 488 conjugate, 53-9760-82 and eFluor 570 conjugate, 41-9760-82, eBioscience/Invitrogen), type I collagen (clone E8F4L, rabbit IgG, 72026, Cell Signaling Technology), type III collagen (rabbit IgG, 22734-1-AP, Proteintech), type IV collagen (clone 1042, mouse IgG2b, Alexa Fluor 488 conjugate, 53-9871-82, eBioscience/Invitrogen), β1 integrin (clone EPR16895, rabbit IgG, ab179471, Abcam), CD235a/glycophorin A (goat IgG, LS-B3010, LSBio and clone HIR2 (GA-R2), mouse IgG2b, biotin conjugate, 13-9987-82, eBioscience/Invitrogen), albumin (rabbit IgG, 16475-1-AP, Proteintech), lysosomal-associated membrane protein 2 (clone H4B4, mouse IgG1, ab25631, Abcam), CD42b/glycoprotein Ib alpha (rabbit IgG, 12860-1-AP, Proteintech), fibrinogen (goat IgG, fluorescein conjugate, 55169, Cappel), and CD45/leukocyte common antigen (goat IgG, LS-B14248, LSBio). Secondary labelling of unconjugated primary antibodies was performed with Alexa Fluor 488, 555, 568, 647, 750, and DyLight 755 conjugated IgG, and Alexa Fluor 647 conjugated streptavidin (Invitrogen). Elastin was labelled with Alexa Fluor 633 hydrazide (A30634, Invitrogen) and DNA with DAPI (D1306, Invitrogen). Sections were mounted with ProLong Gold Antifade (P36984, Thermo Fisher Scientific). Images were acquired using a Stellaris 8 Falcon confocal microscope with LAS X software (Leica).

## Acknowledgments

AC, AHMH, GT, and RA designed the study. AC and AHMH conducted experiments and acquired data. GT and RA contributed specimens. AC, AHMH, JDH, GT, and RA analyzed and interpreted the results. GT and RA supervised the work. AC, AHMH, JDH, GT, and RA wrote and edited the manuscript. The two first authors made equal contributions to the work and the two last authors equally shared supervision of the work.

## List of Abbreviations

ECM: extracellular matrix
RBC: red blood cell
SMC: smooth muscle cell

