## Supplemental Materials for "Hydraulic fracturing-induced delamination and extravasation extends medial damage beyond the false lumen in aortic dissection"

**Supplemental materials:** 2 tables, 7 figures

**Correspondence:** George Tellides, 10 Amistad Street, 337B, New Haven, CT 06520, USA. Phone: +1-203-737-2298, or Roland Assi, 330 Cedar Street, BB 225, New Haven, CT 06510, USA. Phone: +1-203-785-5000,

**Supplemental Table 1: Characteristics of subjects with nondissected aortas***

| | **Demographics**<br>Age, sex, race, ethnicity | **Ascending aorta**<br>Diameter, disease | **Cardiovascular**<br>Risk factors | **Aortic valve**<br>Function, leaflets |
|---|---|---|---|---|
| 1. | 28 yr, male, white, non-Hispanic | 2.7 cm, no disease | Smoker, IVDU | No AI, tricuspid |
| 2. | 40 yr, male, white, Hispanic | 3.6 cm, no disease | Smoker | No AI, tricuspid |
| 3. | 71 yr, female, white, non-Hispanic | 2.8 cm, no disease | Smoker | No AI, tricuspid |
| 4. | 53 yr, male, white, non-Hispanic | 3.0 cm, no disease | Smoker | No AI, tricuspid |
| 5. | 39 yr, male, black, non-Hispanic | 2.8 cm, no disease | Smoker | No AI, tricuspid |
| 6. | 32 yr, male, white, Hispanic | 3.4 cm, no disease | Smoker, cocaine use | No AI, tricuspid |

*Ascending aorta specimens were procured with IRB approval from organ donors without known aortic disease (*n* = 6). Information was obtained from the electronic medical records and intra-operative observations. AI: aortic insufficiency, IVDU: intravenous drug use.

**Supplemental Table 2: Characteristics of subjects with aortic dissection***

| | Demographics<br>Age, sex, race, and ethnicity | Ascending aorta<br>Diameter and associated findings | Aortic dissection<br>Extent and tear location | Clinical presentation<br>Symptoms, onset to XCT, and operative findings | Cardiovascular<br>Diseases and risk factors | Aortic valve<br>Function and leaflet number | Surgery<br>Aortic operation and associated procedures |
|---|---|---|---|---|---|---|---|
| 1. | 54 yr, male, white non-Hispanic | 5.0 cm | Dissection from root to infrarenal aorta, tear in root | Chest pain, shock, 3 hr, tamponade | Alcohol abuse | Severe AI, tricuspid | Root/ascending/hemiarch replacement and antegrade TEVAR |
| 2. | 88 yr, female, white non-Hispanic | 5.4 cm aberrant RSA | Dissection of ascending aorta and arch, chronic dissection distally, multiple tears of ascending aorta | Syncope, hypotension, 4 hr, tamponade | HTN, HLD, chronic type B aortic dissection | Mild AI, tricuspid | Root/ascending/hemiarch replacement |
| 3. | 63 yr, male, white non-Hispanic | 4.2 cm | Dissection from ascending to iliac arteries, tear in proximal arch | Chest, back, and abdominal pain, 20 hr | HTN, obesity, family history of aortic dissection | No AI, tricuspid | Ascending/hemiarch replacement, aortic valve repair, and antegrade TEVAR |
| 4. | 72 yr, female, white non-Hispanic | 4.8 cm | Dissection of ascending aorta, arch, and descending aorta, tear in lesser curve of ascending aorta | Chest pain, 7 hr, hemopericardium | HTN, HLD | Mild AI, tricuspid | Ascending/hemiarch replacement and aortic valve repair |
| 5. | 66 yr, male, white non-Hispanic | 5.3 cm | Dissection of ascending and arch, tears in greater curve above STJ and proximal descending aorta | Chest pain, 7 hr, tamponade | HTN, obesity | No AI, tricuspid | Ascending/zone 2 arch replacement, antegrade TEVAR, and aortic valve repair |
| 6. | 78 yr, female, Pacific Islander, Hispanic | 4.1 cm | Dissection from root to renal arteries, tear above right coronary artery | Chest pain, cardiac arrest, 5 hr, tamponade | HTN, HLD, DM, CAD, Afib, CKD, obesity | No AI , tricuspid | Root/ascending/hemiarch replacement and repair of coronary arteries |
| 7. | 81 yr, male, white non-Hispanic | 4.8 cm | Dissection of root, ascending aorta, and proximal arch, tear in proximal arch | Chest and neck pain, 24 hr, tamponade | HTN, CKD, obesity | Mild AI, tricuspid | Ascending/zone 2 arch replacement |
| 8. | 50 yr, male, white non-Hispanic | 4.6 cm | Dissection from root to abdominal aorta, tear in ascending aorta | Jaw, chest, and back pain, 5 hr | HTN, smoker | Mild AI, tricuspid | Ascending/hemiarch replacement and antegrade TEVAR |
| 9. | 77 yr, female, white non-Hispanic | 4.2 cm | Dissection of root to mid-arch, tear in lesser curve of ascending aorta | Chest pain, 23 hr | HTN, HLD, MI, CAD, COPD | Moderate AI, tricuspid | Ascending/hemiarch replacement and aortic valve repair |
| 10. | 45 yr, male, white non-Hispanic | 8.0 cm 6.4 cm root | Dissection of ascending aorta and arch, tear of proximal ascending aorta | Chest pain, shock, 5 hr, hemopericardium | None | Moderate AI, bicuspid | Root/ascending/hemiarch replacement and repair of right coronary artery |
| 11. | 59 yr, female, white non-Hispanic | 4.0 cm | Dissection from root to descending thoracic, tear in mid-ascending aorta | Chest and back pain, 4 hr, tamponade | HTN, HLD, DM | No AI, tricuspid | Root/ascending/hemiarch replacement, bypass graft to right coronary artery, and repair of left main coronary artery |

*Ascending aortic specimens were procured with IRB approval from patients with acute aortic dissection (*n* = 11). Information was obtained from electronic medical records, imaging studies, and intra-operative observations. Duration of aortic dissection is from the onset of symptoms to intra-operative cross-clamp time. XCT: cross-clamp time, AI: aortic insufficiency, TEVAR: thoracic endovascular aortic repair, RSA: right subclavian artery, STJ: sinotubular junction, HTN: hypertension, HLD: hyperlipidemia; DM: diabetes mellitus, CAD: coronary artery disease, MI: prior myocardial infarction, Afib: atrial fibrillation, CKD: chronic kidney disease, COPD: chronic obstructive pulmonary disease.

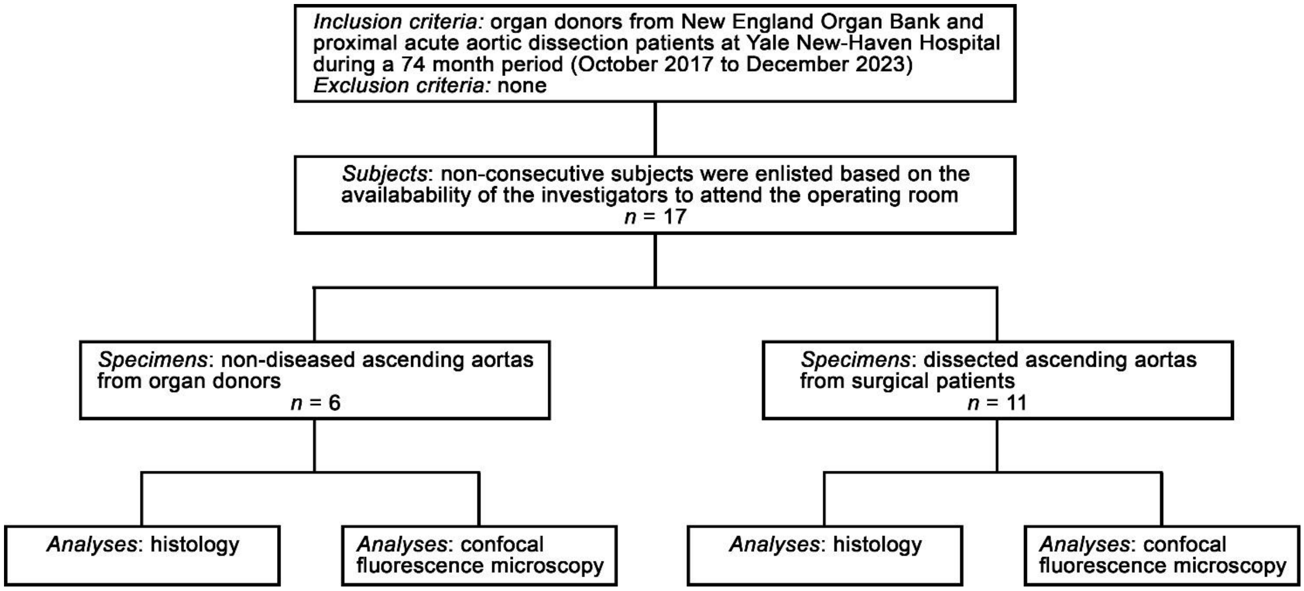


**Supplemental Figure 1: Flow diagram of study design.** Summary of inclusion/exclusion criteria, number of subjects enlisted, and number of specimens analyzed. Histology and confocal fluorescence microscopy were performed in 6 non-diseased aortas from organ donors and 11 acutely dissected aortas from surgical patients. Patients with aortic dissection compared to organ donors without aortic disease did not differ by sex (45.5% vs. 16.7% female, $P$ = 0.33, Fisher's exact test), but were older (66.6 ± 13.8 vs. 43.8 ± 15.8 yr, $P$ = 0.0153, unpaired t-test with Welch's correction) and had larger ascending aortas (4.9 ± 1.1 vs. 3.1 ± 0.4 cm, $P$ = 0.0002, Mann-Whitney test); individual data shown in Supplemental Tables 1 and 2.

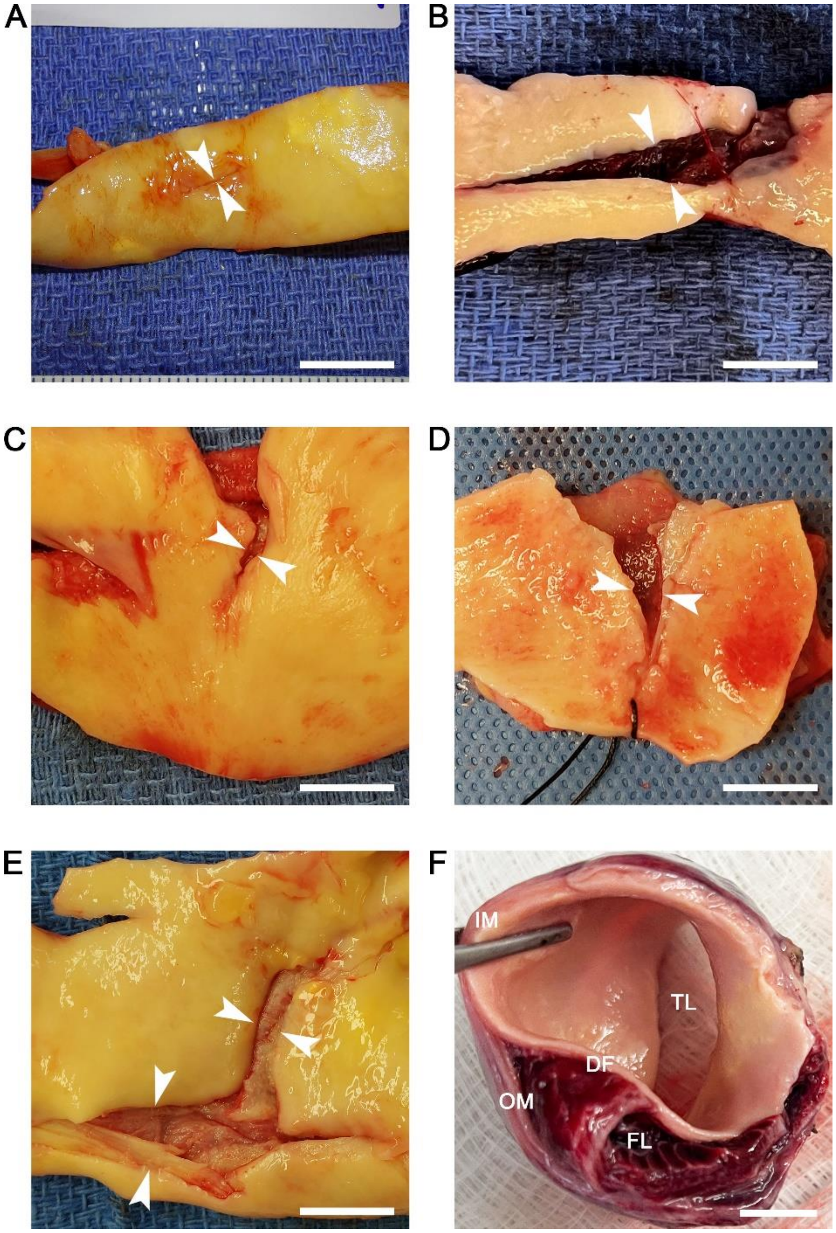


**Supplemental Figure 2: Gross appearance of aortic dissection lesions.** Specimens of proximal ascending aortas excised during surgical repair for dissection were examined in the operating room. (**A**) Transverse entry tear without retraction of edges. (**B**) Transverse entry tear with marked retraction of edges. (**C**) Longitudinal entry tear with mild retraction of edges. (**D**) Longitudinal entry tear with moderate retraction of edges. (**E**) Curvilinear entry tear with marked retraction of edges. (**F**) Transverse section through dissected ascending aorta showing true lumen (TL) and false lumen (FL) filled with thrombus; intact media (IM) that is not dissected bordering the true lumen, dissection flap (DF) consisting of intima and dissected inner media partitioning the true from false lumen, and dissected outer media (OM) with adventitia bordering the outer false lumen. Edges of entry tears are delineated by arrows. Orientation: proximal aspect of aorta at bottom and distal aspect at top (panels A–E), and intact media above and dissected media below (panel F). Scale bars = 1 cm.

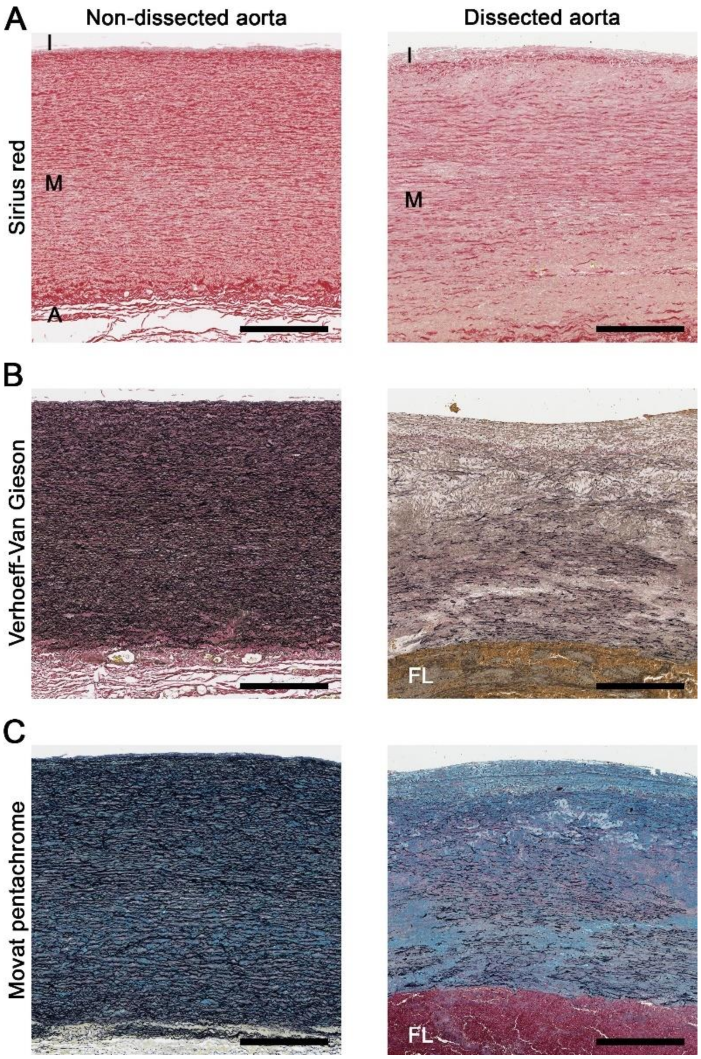


**Supplemental Figure 3: Medial structure in non-dissected and dissected aortas.** Histology of ascending aortas from an organ donor without medial degeneration (left panels) and acute dissection with pre-existing severe medial degeneration (right panels). (**A**) Sirius red stain labeling collagen (red) delineating the intima (I), media (M), and adventitia (A) layers. (**B**) Verhoeff-Van Gieson stain labeling elastin (black) and RBCs (yellow) delineating the false lumen (FL). (**C**) Movat pentachrome stain labeling elastin (black), SMCs and RBCs/thrombus (red), collagen (yellow), and glycosaminoglycans (blue). Orientation: luminal side top, adventitial side bottom. Scale bars = 500 μm.

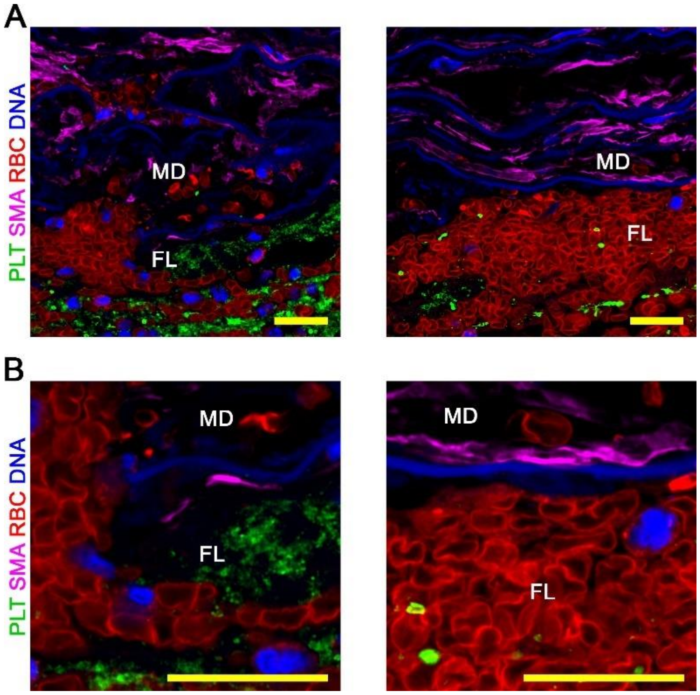


**Supplemental Figure 4: Platelet distribution in aortic dissection lesions.** Confocal fluorescence microscopy images of dissected aortas. (**A**) Lower and (**B**) higher magnification views for CD42b (green) identifying platelets (PLT), smooth muscle α-actin (SMA, magenta) identifying SMCs, CD235a (red) identifying RBCs, and DAPI (blue) labeling nuclei that show numerous platelets within the false lumen (FL) but few in adjacent delaminated laminae filled with RBCs, termed minor delaminations (MD). Elastic lamellae are faintly visible from blue autofluorescence. Orientation: luminal side top, adventitial side bottom. Scale bars = 20 μm.

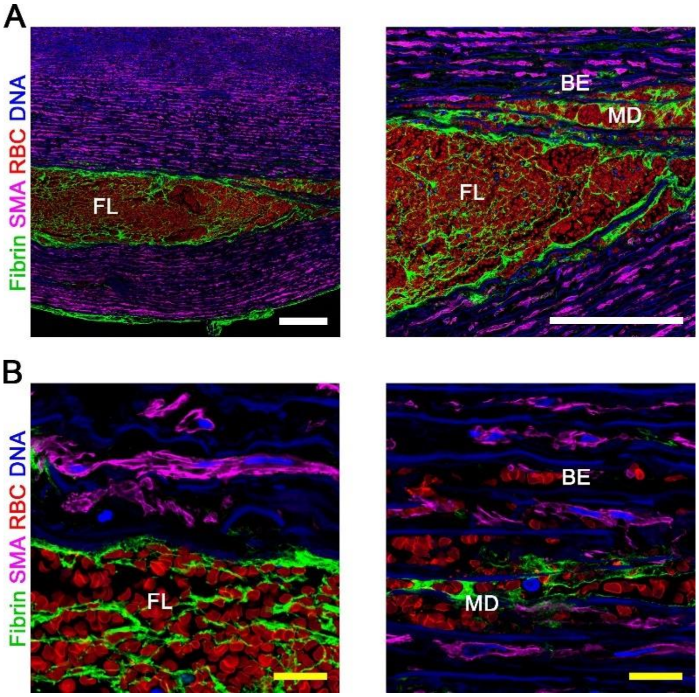


**Supplemental Figure 5: Fibrin distribution in aortic dissection lesions.** Confocal fluorescence microscopy images of dissected aortas. (**A**) Lower and (**B**) higher magnification views for fibrin (green) delineating thrombus, smooth muscle α-actin (SMA, magenta) identifying SMCs, CD235a (red) identifying RBCs, and DAPI (blue) labeling nuclei that show fibrin deposition within the false lumen (FL) and minor delaminations (MD) but rarely in non-delaminated laminae with RBCs, termed blood extravasation (BE). Elastic lamellae are faintly visible from blue autofluorescence. Orientation: luminal side top, adventitial side bottom. Scale bars: white = 250 μm, yellow = 20 μm.

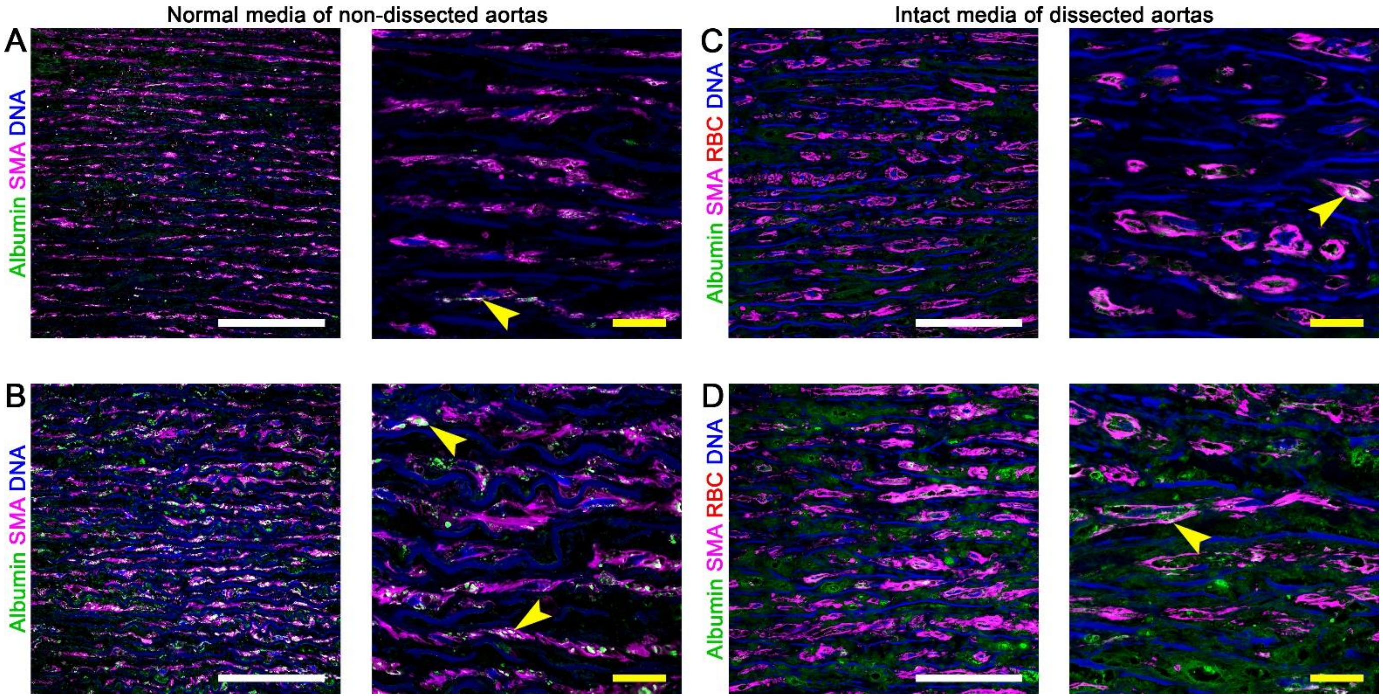


**Supplemental Figure 6: Albumin distribution in non-dissected and dissected aortas.** Confocal fluorescence microscopy images of aortic media. Lower (left panels) and higher (right panels) magnification views for albumin (green), smooth muscle α-actin (SMA, magenta) identifying SMCs, CD235a (red) identifying RBCs, and DAPI (blue) labeling nuclei show a spectrum of albumin accumulation from (**A**) minimal to (**B**) moderate in normal media of non-dissected aortas, and (**C**) minimal to (**D**) moderate in the intact media of dissected aortas. Overlay of intracellular albumin (green) and cytoplasmic smooth muscle α-actin (magenta) appears white (arrows). Elastic lamellae are faintly visible from blue autofluorescence. Orientation: luminal side top, adventitial side bottom. Scale bars: white = 100 μm, yellow = 20 μm.

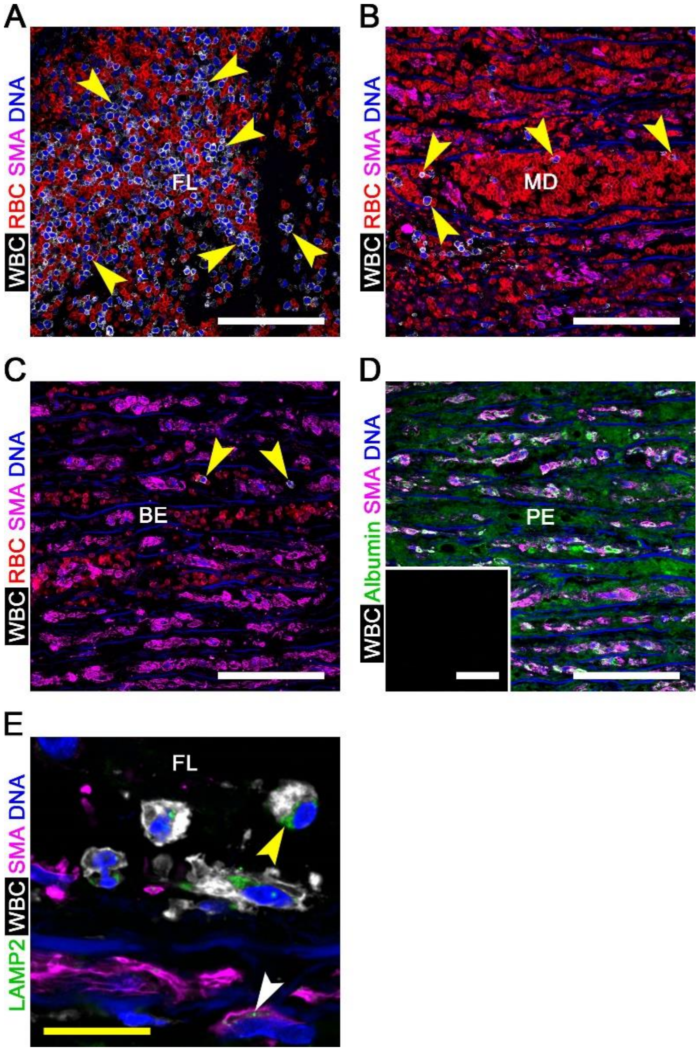


**Supplemental Figure 7: Leukocyte distribution in aortic dissection lesions.** Confocal fluorescence microscopy images of dissected aortas. CD45 (white) identifying leukocytes (WBC), CD235a (red) identifying RBCs, smooth muscle α-actin (SMA, magenta) identifying SMCs, and DAPI (blue) labeling nuclei showing (**A**) numerous leukocytes (arrows) in false lumen (FL), (**B**) frequent leukocytes (arrows) in minor delaminations (MD), and (**C**) infrequent leukocytes (arrows) in lesion areas with blood extravasation (BE). (**D**) Additional staining for albumin (green) showing absence of leukocytes in lesion areas with plasma extravasation (PE); fluorescence signal for CD45 was not detected (inset) though white color within SMCs results from overlay of magenta (SMA) and green (intracellular albumin) pseudocolors. (**E**) Alternatively, delineation of endolysosomes by lysosomal-associated membrane protein 2 (LAMP2, green), leukocytes by CD45 (white), SMCs by smooth muscle α-actin (magenta), and nuclei by DAPI (blue) verifies readily detectable organelles in mononuclear leukocytes (yellow arrow) but few in SMCs (white arrow). Elastic lamellae are faintly visible from blue autofluorescence. Orientation: luminal side top, adventitial side bottom. Scale bars: white = 100 μm, yellow = 20 μm.
